# Reconstitution of prenyltransferase activity on nanodiscs by components of the rubber synthesis machinery of the Para rubber tree and guayule

**DOI:** 10.1101/2021.04.29.441905

**Authors:** Fu Kuroiwa, Akira Nishino, Yasuko Mandal, Miki Suenaga-Hiromori, Kakeru Suzuki, Yukie Takani, Yukino Miyagi-Inoue, Haruhiko Yamaguchi, Satoshi Yamashita, Seiji Takahashi, Yuzuru Tozawa

**Author notes:** Corresponding author: Yuzuru Tozawa, Graduate School of Science and Engineering, Saitama University, 255 Shimo-Okubo, Sakura-ku, Saitama, Saitama 338-8570, Japan.

## Abstract

Prenyltransferases mediate the biosynthesis of various types of polyisoprene compound in living organisms. Natural rubber (NR) of the Para rubber tree (*Hevea brasiliensis*) is synthesized as a result of prenyltransferase activity, with the proteins HRT1, HRT2, and HRBP having been identified as candidate components of the rubber biosynthetic machinery. To clarify the contribution of these proteins to prenyltransferase activity, we established a cell-free translation system for nanodisc-based protein reconstitution and measured the enzyme activity of the protein-nanodisc complexes. Cell-free synthesis of HRT1, HRT2, and HRBP in the presence of asolectin nanodiscs revealed that all three proteins were membrane associated. A complex of HRT1 and HRBP formed as a result of co-expression of the two proteins in the presence of nanodiscs manifested marked polyisoprene synthesis activity, whereas neither HRT1, HRT2, or HRBP alone nor a complex of HRT2 and HRBP exhibited such activity. Similar analysis of guayule (*Parthenium argentatum*) proteins revealed that three HRT1 homologs (CPT1–3) manifested prenyltransferease activity only if co-expressed with the homolog of HRBP (CBP). Our results thus indicate that the core prenyltransferase of the rubber biosynthetic machinery of both the Para rubber tree and guayule is formed by the assembly of heterologous subunits (HRT1 and HRBP in the former species).

## 1. Introduction

Natural rubber (NR) is an industrially important product whose supply has been mainly dependent on the Para rubber tree (*Hevea brasiliensis*). The unique physical properties of NR allow the manufacture of a variety of polymer products. However, the production of NR from the rubber tree is largely limited to Southeast Asia, and a stable supply of NR to support expansion of demand is not guaranteed. NR has therefore been considered a strategically important natural product in many countries. Plants such as guayule (*Parthenium argentatum*) and Russian dandelion (*Taraxacum kok-saghyz*) have been identified as potential alternative sources for the production of NR, and the NR biosynthetic systems of these species as well as of the Para rubber tree have been investigated [1–6].

NR is an isoprenoid with a backbone structure comprising a polymerized C_5_H_8_ isoprene unit with a *cis*-1,4 configuration. This backbone is also present in dolichols, which function as carriers during protein glycosylation in eukaryotes. However, the mechanism underlying the marked polymerization characteristic of NR biosynthesis, which results in the production of polyisoprene with a molecular size of ~1 MDa, has remained to be elucidated. Our collaborative group previously isolated two *cis*-prenyltransferase (CPT) homologs, designated *Hevea brasiliensis* rubber transferase (HRT) 1 and HRT2, from *H. brasiliensis*, with these proteins being predominantly localized to rubber particles [7]. Expression of HRT1 or HRT2 in a yeast ∆*rer2* mutant, which lacks its major CPT activity, revealed that each of these proteins possesses enzymatic activity capable of producing C_80_–C_100_ polyisoprenoids [8]. Cell-free synthesis of HRT1 in the presence of detergent-washed rubber particles derived from *H. brasiliensis* allowed reconstitution of NR biosynthetic activity in vitro [9]. An *H. brasiliensis* homolog of the mammalian CPT subunit NgBR that interacts with HRT1 and rubber elongation factor (REF) was also identified and designated HRT1-REF bridging protein (HRBP). The NR biosynthesis machinery was thus proposed to include HRT1 and HRBP as well as the key factors REF and small rubber particle protein (SRPP) [9].

Cell-free systems provide an effective tool for the production of membrane proteins [10,11], with various modified such systems having been described [12,13]. A cell-free system supplemented with nanodiscs has been developed for the production of *Escherichia coli* membrane proteins with functional folds [14]. Nanodiscs are disclike forms of lipid-bilayer membranes surrounded by a membrane scaffold protein (MSP) [15]. To gain insight into the core enzyme complex of the rubber biosynthetic machinery, we have now established a nanodisc-supplemented cell-free system for reconstitution of prenyltransferase activity (Fig. S1) and have characterized the enzyme activities of purified protein-nanodisc complexes. We found that the combination of HRT1 and HRBP reconstitutes prenyltransferase activity on nanodiscs. We also show that the guayule homologs of these Para rubber tree proteins likewise reconstitute an active prenyltransferase enzyme on nanodiscs.

## 2. Materials and Methods

### 2.1. Nanodisc preparation

A synthetic gene encoding the MSP derivative MSP1E3D1 with an NH_2_-terminal His_6_ tag [15] was obtained from Eurofins Genomics. Expression and purification of MSP1E3D1 were performed as described previously [14], with minor modifications. After bacterial cell culture, cells isolated by centrifugation were resuspended in a binding buffer comprising 50 mM Tris-HCl (pH 7.5), 300 mM NaCl, and 20 mM imidazole. The cells were then subjected to extraction by ultrasonic treatment, the extract was centrifuged at 20,000 × *g* for 30 min at 4°C, and the resultant supernatant was mixed with 2 ml of Ni-NTA agarose resin suspension (Qiagen) and incubated on a rotary shaker for 60 min at 4°C before transfer to a Poly-Prep Chromatography Column (Bio-Rad). The column bed was washed with 40 ml of a wash buffer containing 50 mM Tris-HCl (pH 7.5), 300 mM NaCl, and 50 mM imidazole before elution with a solution containing 50 mM Tris-HCl (pH 7.5), 300 mM NaCl, and 250 mM imidazole. The eluted fractions were collected in 1.5-ml tubes, and those containing MSP1E3D1 were combined and subjected to dialysis (molecular size cutoff of 12 to 14 kDa) for 2 days at 25°C with a 500-fold volume of a solution containing 50 mM Tris-HCl (pH 7.5) and 300 mM NaCl, with four changes of the dialysis buffer.

For nanodisc reconstitution, the MSP1E3D1 solution was mixed with an equal volume of 600 mM sodium cholate and agitated gently for 20 min at 25°C and was then combined with other components to form a mixture containing 10 mM Tris-HCl (pH 8.0), 100 mM NaCl, 162 mM sodium cholate, 40 μM MSP1E3D1, and 2 mM asolectin (Sigma-Aldrich). The mixture was agitated gently for 60 min at 25°C and then subjected to dialysis (molecular size cutoff of 12 to 14 kDa) for 2 days at 25°C with a 500-fold volume of ND buffer comprising 10 mM Tris-HCl (pH 8.0) and 100 mM NaCl, with four changes of the latter buffer. The mixture was centrifuged at 20,400 × *g* for 30 min at 4°C, and the resulting supernatant was applied to a gel-filtration column, Superdex 200 Increase 10/300 GL (Cytiva). Chromatography was performed with ND buffer at a flow rate of 0.5 ml/min and was monitored by measurement of absorbance at 280 nm, and the peak fractions corresponding to the assembled nanodiscs were collected. The eluate containing the nanodiscs was concentrated to 1 ml with the use of an Amicon Ultra-15 10K device (Merck Millipore) and then mixed with 10 ml of translation buffer containing 30 mM HEPES-KOH (pH 7.8) and 100 mM potassium acetate before adjustment of the nanodisc concentration to 200 μM.

### 2.2. Cell-free protein synthesis

Codon usage for nucleotide sequences of protein-coding regions were optimized for *Triticum aestivum* (wheat), and the DNA molecules were synthesized at Eurofins Genomics. GenBank accession numbers for the modified cDNAs are LC626733, LC626734, and LC626735 for *HRT1*, *HRT2*, and *HRBP*, respectively, of *H. brasiliensis*; LC626736, LC626737, LC626738, and LC626739 for *CPT1*, *CPT2*, *CPT3*, and *CBP*, respectively, of *P. argentatum*; and LC626740 for human *β2AR*. The synthesized sequences were cloned into the pYT08 vector [12]. Wheat-germ extract for cell-free protein synthesis (WEPRO 7240 or 7240H) was obtained from CellFree Sciences. Transcription and cell-free protein synthesis were performed as previously described [16], with some modifications. Template mRNAs for cell-free protein synthesis were produced by in vitro transcription with SP6 RNA polymerase. Nucleotide sequences for the recombinant proteins were optimized for the wheat-germ cell-free system and cloned into the pYT08 vector. Plasmids were amplified in *E. coli*, isolated with the use of a Plasmid Maxi Kit (Qiagen), and then subjected to in vitro transcription in a transcription mixture of 100 μl containing 10 μg of plasmid as template DNA, 40 mM Tris-HCl (pH 7.5), 6 mM MgCl_2_, 2 mM spermidine, 10 mM dithiothreitol, 0.01% BSA, 3 mM of each nucleoside triphosphate, RNase inhibitor (0.8 U/μl) (Promega), and SP6 RNA polymerase (0.8 U/μl) (Promega). The transcription mixture was gently mixed by pipetting and incubated at 37°C for 3 h in a water bath. The mixture was then centrifuged at 20,000 × *g* for 1 min, and the resulting supernatant was transferred to a new microtube, mixed with 13 μl of 7.5 M ammonium acetate and 250 μl of 100% ethanol, and incubated at –20°C for 15 min. The synthesized mRNAs were isolated by centrifugation at 20,000 × *g* for 20 min at 4°C, and the resulting pellet was washed with 800 μl of 70% ethanol, isolated again by centrifugation, allowed to dry in air, and then dissolved in 21 μl of deionized water. The translation mixture (total volume of 50 μl) contained 30 mM HEPES-KOH (pH 7.8), 100 mM potassium acetate, 2.7 mM magnesium acetate, 0.4 mM spermidine, 2.5 mM dithiothreitol, 0.3 mM of each amino acid, 1.2 mM ATP, 0.25 mM GTP, 16 mM creatine phosphate, creatine kinase (0.4 mg/ml), RNase inhibitor (0.8 U/μl), 15 μM asolectin nanodiscs, 15 μl of wheat-germ extract (CellFree Sciences), and 100 μg of mRNA (in the case of co-expression of two different proteins, 50 μg of each mRNA were added). The reaction mixture was placed in a dialysis cup with a molecular size cutoff of 12 kDa (Biotech International) and dialyzed against 1 ml of substrate supply solution containing 30 mM HEPES-KOH (pH 7.8), 100 mM potassium acetate, 2.7 mM magnesium acetate, 0.4 mM spermidine, 2.5 mM dithiothreitol, 0.3 mM of each amino acid, 1.2 mM ATP, 0.25 mM GTP, 16 mM creatine phosphate, and 0.005% NaN_3_. The mixture was then incubated at 16°C for 48 h.

### 2.3. Purification of protein-nanodisc complexes

Protein-nanodisc complexes were purified by immobilized metal affinity chromatography (IMAC) based on the interaction of His_6_-tagged MSP1E3D1 with Ni-NTA agarose. The translation mixture was gently agitated in a microtube for 1 h at 4°C with a fivefold volume of Ni-NTA agarose (Qiagen) that had been equilibrated with a solution containing 50 mM Tris-HCl (pH 7.5) and 300 mM NaCl. The mixture was then transferred to a Poly-Prep Chromatography Column (Bio-Rad), the column was washed with three column-bed volumes of a solution containing 50 mM Tris-HCl (pH 7.5), 300 mM NaCl, and 50 mM imidazole, and the protein-nanodisc complexes were eluted with a solution containing 50 mM Tris-HCl (pH 7.5), 300 mM NaCl, and 250 mM imidazole and then concentrated to ~50 μl by centrifugation in a VIVASPIN 500 tube (Sartorius), during which the buffer was exchanged with 50 mM Tris-HCl (pH 7.5). The protein-nanodisc complexes were denatured by incubation in the presence of 2% SDS and 5% 2-mercaptoethanol for 1 h at 37°C for confirmation of their composition by SDS–polyacrylamide gel electrophoresis (PAGE) on a 12% acrylamide gel. The gel was stained with Coomassie brilliant blue (CBB) (Bio Craft), and the concentration of the purified protein in the samples was estimated relative to that of bovine serum albumin (BSA) standard solutions. Band intensity was calculated with the use of ImageJ software [17] (Fig. S2). Given that the molecular sizes of CPT1 of *P. argentatum* (PaCPT1) and MSP are similar and that their bands overlapped in CBB-stained gels, the ratio of PaCPT1 to MSP was estimated on the basis of the relative ratio of PaCPT2 to MSP in the corresponding complex.

### 2.4. Measurement of prenyltransferase activity

The prenyltransferase activity of protein-nanodisc complexes was measured with a modified version of a previously described method [9]. Purified protein-nanodisc complexes were added to a reaction mixture (final volume of 50 μl) containing 50 mM Tris-HCl (pH 7.5), 2 mM dithiothreitol, 4 mM MgCl_2_, 15 μM farnesyl pyrophosphate, and 125 nCi of [4-^14^C]isopentenyl pyrophosphate (IPP) (Perkin Elmer, 50.6 mCi/mmol) in deionized water. The amounts of protein-nanodisc complexes added to the enzyme assay were adjusted so that 0.5 μg of HRT1, HRT2, PaCPT1, PaCPT2, PaCPT3, or β2AR (negative control) was contained in each reaction mixture. For the protein-nanodisc complexes without HRT or CPT, the amounts were adjusted so that 0.5 μg of HRBP or PaCBP was contained in each reaction mixture. The reaction mixtures were incubated for 20 h at 16°C, after which 100 μl of deionized water saturated with NaCl were added and polyisoprenes with a common chain length shorter than that of NR were extracted with 500 μl of 1-butanol that had been saturated with NaCl-saturated water. The NR-length polyisoprenes in the remaining aqueous layer were extracted with 500 μl of toluene/hexane (1:1, v/v). The amount of radioactivity in the extracts was measured with a liquid scintillation counter. The enzyme assay was performed in triplicate, and background counts were measured for reaction mixtures without protein-nanodisc complexes according to the same procedure. The background value was subtracted from each experimental value to determine the counts for [^14^C]IPP incorporation.

### 2.5. Reversed-phase TLC analysis

Analysis of chain length for polyisoprenes extracted by 1-butanol was performed as previously described [8]. The extracts were hydrolyzed with acid phosphatase (Sigma) to produce alcohols, which were then extracted with pentane [18] for analysis by reversed-phase thin-layer chromatography (TLC) with HPTLC Silica gel 60 RP-18 (Merck) and an acetone/water (39:1, v/v) solvent system. The [^14^C]IPP polymers spread on the TLC plate were detected with a BAS1000 Mac Bioimage Analyzer (Fujifilm). The chain length of the reaction products was estimated with the use of carbon number–defined *Z,E*-mixed polyprenols (C_55_, C_60_, C_85_, and C_90_) as authentic standards, as described previously [8].

## 3. Results

### 3.1. Cell-free synthesis and enzyme assay for HRT1, HRT2, and HRBP proteins

To investigate the functions of HRT1, HRT2, and HRBP, we synthesized each protein with a wheat-germ cell-free translation system. Given that HRBP is thought to function cooperatively with HRT1 or HRT2 [9], we also attempted co-translation of the combinations of HRBP and either HRT protein with the cell-free system. The human β2-adrenergic receptor (β2AR) was synthesized as a control [19]. The cDNAs for HRT1, HRT2, HRBP, and β2AR were cloned into the cell-free expression plasmid pYT08, and the resultant plasmids were subjected to transcription and subsequent cell-free protein synthesis in the presence of preassembled asolectin nanodiscs (Fig. S1). The resulting protein-nanodisc complexes were isolated by IMAC and analyzed by SDS-PAGE and CBB staining (Fig. 1A). The molecular masses of HRT1, HRT2, HRBP, β2AR, and MSP (a component of the nanodiscs) derived from their electrophoretic mobilities were essentially consistent with the predicted values of 33.1, 32.7, 29.4, 46.5, and 32.6 kDa, respectively. The results thus indicated that each synthesized protein was associated with the nanodiscs purified from the translation reaction mixture, revealing that HRT1, HRT2, and HRBP behave as membrane proteins.

**Fig. 1.**
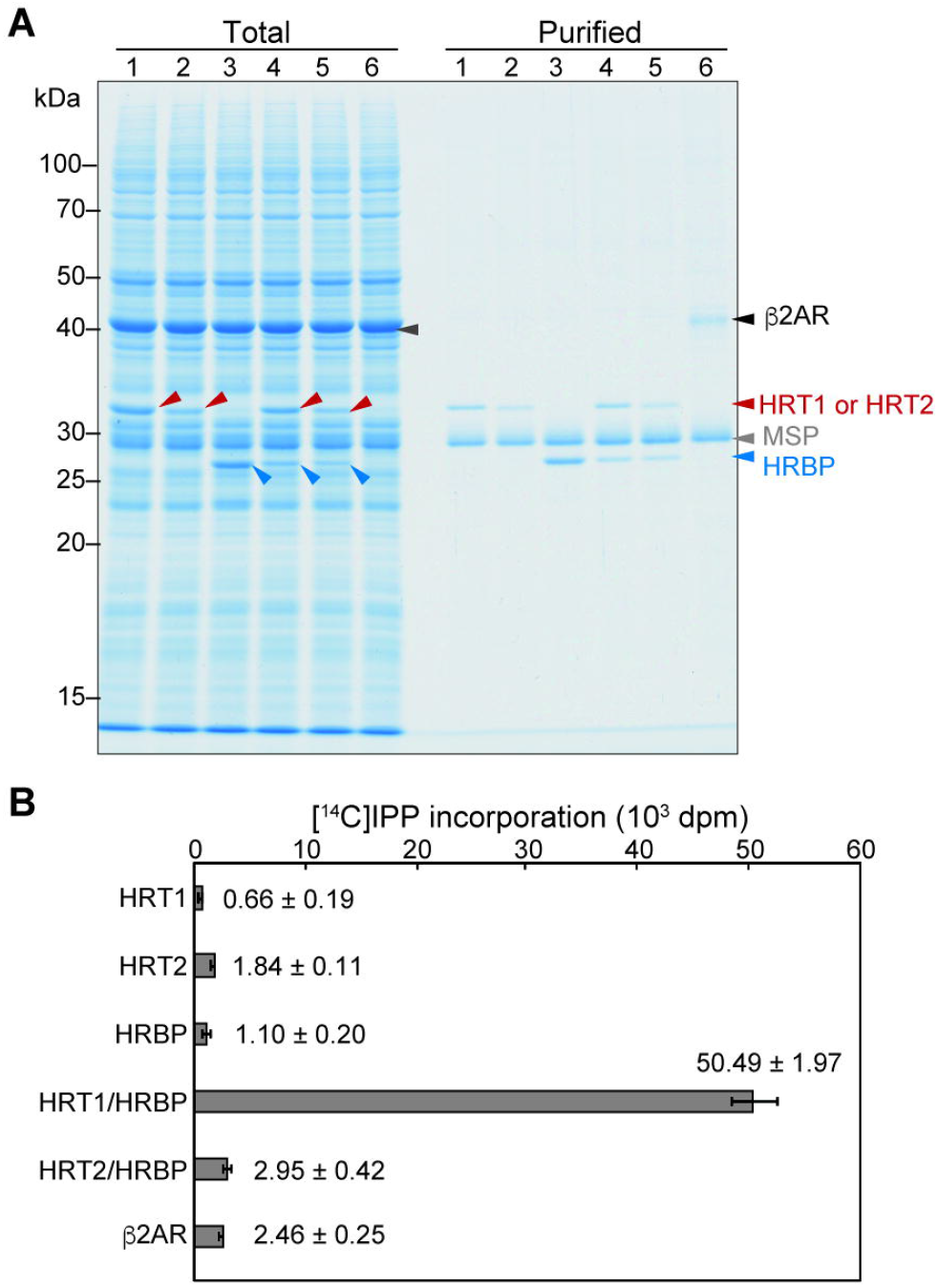
Preparation and characterization of protein-nanodisc complexes. (**A**) HRT1, HRT2, HRBP, HRT1 plus HRBP, HRT2 plus HRBP, and β2AR (lanes 1–6, respectively) were synthesized with a wheat-germ cell-free system in the presence of nanodiscs. The protein-nanodisc complexes were purified by Ni-NTA column chromatography and, together with a portion of the translation reaction mixture corresponding to 1% of the input for purification (Total), were subjected to SDS-PAGE and staining with CBB. Arrowheads indicate HRT1 or HRT2 (red), HRBP (blue), β2AR (black), and MSP (gray). The positions of molecular size markers are indicated on the left. (**B**) Polyisoprene synthesis activity assay for the purified protein-nanodisc complexes in (**A**). The incorporation of [^14^C]IPP into macromolecules extracted with 1-butanol was measured. Data are means ± SD from three independent experiments.

We next assessed each protein-nanodisc complex for prenyltransferase activity. The isolated complexes were thus assayed for polyisoprene synthesis activity with [^14^C]IPP. Measurement of [^14^C]IPP incorporation into macromolecules extracted from the reaction mixture with 1-butanol (Fig. 1B). The nanodiscs containing HRT1, HRT2, or HRBP alone or those containing both HRT2 and HRBP did not catalyze incorporation of [^14^C]IPP monomer into extracted macromolecules. In contrast, the nanodiscs containing both HRT1 and HRBP mediated marked incorporation of [^14^C]IPP into macromolecules (Fig. 1B). On the other hand, toluene/hexane extracts of none of the various reaction mixtures revealed incorporation of [^14^C]IPP into macromolecules (data not shown). Polyisoprenes with the size of NR were thus not synthesized by the protein products associated with nanodiscs in our in vitro system.

### 3.2. Effect of sequential expression of HRT1 and HRBP on enzymatic activity

Our results showed that co-expression of HRT1 and HRBP in the presence of nanodiscs reconstituted prenyltransferase activity in vitro. We next examined the possible effect of sequential expression of the two proteins on enzyme activity. HRT1-nanodisc and HRBP-nanodisc complexes were each purified and added to the reaction mixture for cell-free protein synthesis of HRBP and HRT1, respectively (Fig. 2A). The nanodiscs were then again purified and analyzed by SDS-PAGE (Fig. 2B), which revealed that each expression sequence resulted in the association of HRT1 and HRBP with nanodiscs. Each of the purified HRT1/HRBP-nanodisc complexes was also found to possess prenyltransferase activity in vitro (Fig. 2C). TLC analysis revealed that the patterns of polyisoprene elongation were also similar for the HRT1/HRBP-nanodisc complexes produced by co-expression or sequential expression of the two proteins (Fig. 2D). On the other hand, simple mixture of HRT1-nanodisc and HRBP-nanodisc complexes did not result in substantial reconstitution of prenyltransferase activity (Fig. 2C, D). These results thus suggested that reconstitution of prenyltransferase activity requires direct contact between HRT1 and HRBP on the surface of nanodiscs, with the order of HRT1 and HRBP protein synthesis in the cell-free system not having a substantial effect on enzyme activity.

**Fig. 2.**
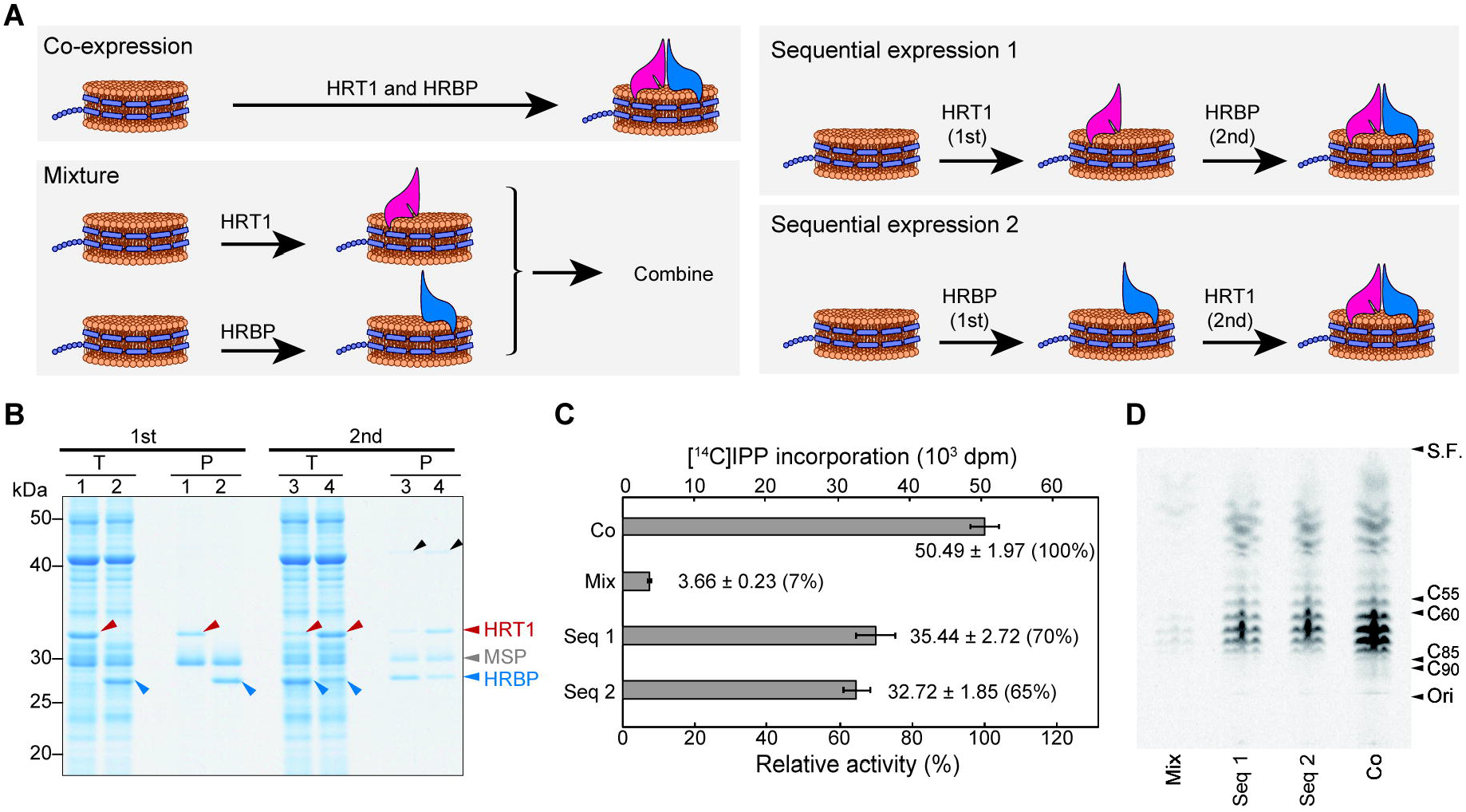
Sequential expression and characterization of HRT1/HRBP-nanodisc complexes. (**A**) Schematic representation of protein expression protocols. The mRNAs for HRT1 and HRBP were translated together in the presence of nanodiscs (Co-expression); purified HRT1-nanodisc and HRBP-nanodisc complexes were mixed (Mixture); or HRBP or HRT1 was synthesized in the presence of HRT1-nanodisc or HRBP-nanodisc complexes, respectively (Sequential expression 1 and 2, respectively). (**B**) Purified protein-nanodisc complexes (P) as well as a portion of the translation reaction mixture corresponding to 1% of the input for purification (T) were subjected to SDS-PAGE and stained with CBB. 1st and 2nd indicate the first and second expression, respectively. Lanes 1 to 4 correspond to expression of HRT1 in the presence of empty nanodiscs, expression of HRBP in the presence of empty nanodiscs, expression of HRBP in the presence of HRT1-nanodisc complexes, and expression of HRT1 in the presence of HRBP-nanodisc complexes, respectively. Arrowheads indicate HRT1 (red), HRBP (blue), and MSP (gray) as well as a co-purified protein from the wheat-germ extract (black). (**C**) Polyisoprene synthesis activity assay for the purified protein-nanodisc complexes. Co, co-expression; Mix, mixture; Seq 1, sequential expression 1; Seq 2, sequential expression 2. The incorporation of [^14^C]IPP into macromolecules extracted with 1-butanol was measured. Data are means ± SD from three independent experiments. (**D**) Analysis of chain length for the ^14^C-labeled polyisoprenes extracted by 1-butanol was performed by TLC and autoradiography. The positions of the origin (Ori), solvent front (S.F.), and authentic standards are indicated on the right side.

### 3.3. Characterization of guayule CPT and CBP proteins

Three HRT1 homologs (CPT1, CPT2, and CPT3) and one HRBP homolog (CPT binding protein, or CBP) have been identified for *P. argentatum* [5]. We therefore next prepared protein-nanodisc complexes for these guayule proteins and assayed them for prenyltransferase activity in vitro. SDS-PAGE analysis of the purified protein-nanodisc complexes indicated that the electrophoretic mobility of PaCPT1 was similar to that of MSP (Fig. 3A), with co-migration of the two proteins being confirmed by radioisotope labeling of PaCPT1 (Fig. S3). The prenyltransferase activity assay (Fig. 3B) and TLC analysis (Fig. 3C) revealed that [^14^C]IPP incorporation into macromolecules extracted by 1-butanol was catalyzed by PaCPT1/PaCBP, PaCPT2/PaCBP, and PaCPT3/PaCBP complexes but not by the individual guayule proteins. TLC analysis of toluene/hexane extracts did not show substantial incorporation of [^14^C]IPP into macromolecules for any of the enzyme assay mixtures, indicating that products corresponding to rubber-size polyisoprenes were not synthesized (data not shown).

**Fig. 3.**
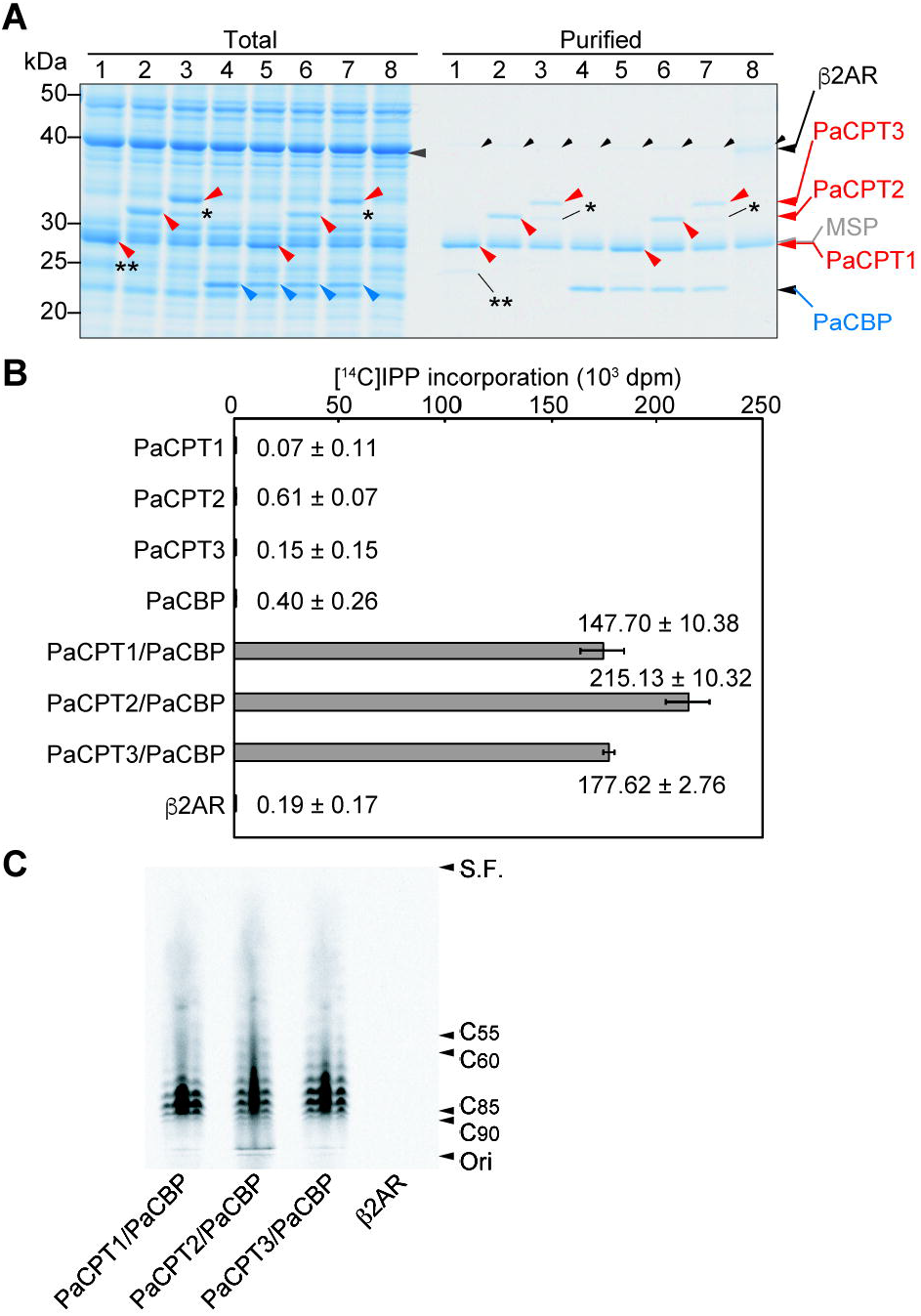
Characterization of guayule homologs of HRT1 and HRBP. (**A**) The guayule proteins PaCPT1, PaCPT2, PaCPT3, and PaCBP were synthesized with a wheat-germ cell-free system in the presence of nanodiscs either separately (lanes 1–4, respectively) or together in the combinations PaCPT1/PaCBP, PaCPT2/PaCBP, or PaCPT3/PaCBP (lanes 5–7, respectively). The protein-nanodisc complexes were then purified and analyzed by SDS-PAGE and CBB staining. β2AR was similarly synthesized and purified as a control (lane 8). A portion of each translation reaction mixture corresponding to 1% of the input for purification (Total) was also analyzed. Arrowheads indicate PaCPT1, PaCPT2, or PaCPT3 (red); PaCBP (blue); β2AR (large black); MSP (gray); or a wheat-germ protein co-purified by Ni-NTA column chromatography (small black). Single and double asterisks indicate a truncated form of PaCPT3 translated from an internal initiation codon (Fig. S4) and an uncharacterized translation product for PaCPT1 mRNA, respectively. (**B**) Polyisoprene synthesis assay for the purified protein-nanodisc complexes in (**A**). The incorporation of [^14^C]IPP into macromolecules extracted with 1-butanol was measured. Data are means ± SD from three independent experiments. (**C**) Analysis of chain length for the ^14^C-labeled polyisoprenes extracted by 1-butanol was performed by TLC and autoradiography.

In the preparation of protein-nanodisc complexes, we observed two distinct bands for the PaCPT3 translation reaction mixture: a larger, major band and smaller, fainter band (Fig. 3A, Fig. S3). The mRNA sequence for PaCPT3 contains an alternative initiation codon that is located downstream of the first AUG and which corresponds to the position of the initiation codon for PaCPT2 (Fig. S4A). We therefore constructed an expression vector for the NH_2_-terminally truncated form of PaCPT3 (PaCPT3_dN15) and prepared a PaCPT3_dN15–nanodisc complex. The electrophoretic mobility of PaCPT3_dN15 matched that of the smaller molecule observed during preparation of the PaCPT3-nanodisc complex (Fig. S4B), and the PaCPT3_dN15/CBP complex manifested prenyltransferase activity similar to that of PaCPT3/PaCBP (Fig. S4C, D). The 15 amino acids at the NH_2_-terminus of PaCPT3 therefore do not appear to affect enzymatic function.

Finally, we investigated the interspecies compatibility of CPT catalytic subunits (HRT1 or PaCPT2) and their associated proteins (HRBP or PaCBP) with regard to reconstitution of enzyme activity with nanodiscs (Fig. 4A). The catalytic activity of PaCPT2 was supported by HRBP, although not to the same extent as it was by PaCBP, whereas that of HRT1 was not supported by PaCBP (Fig. 4B, C). PaCPT2/PaCBP appeared to possess a greater prenyltransferase activity than did HRT1/HRBP, which might account in part for the activity observed with PaCPT2/HRBP.

**Fig. 4.**
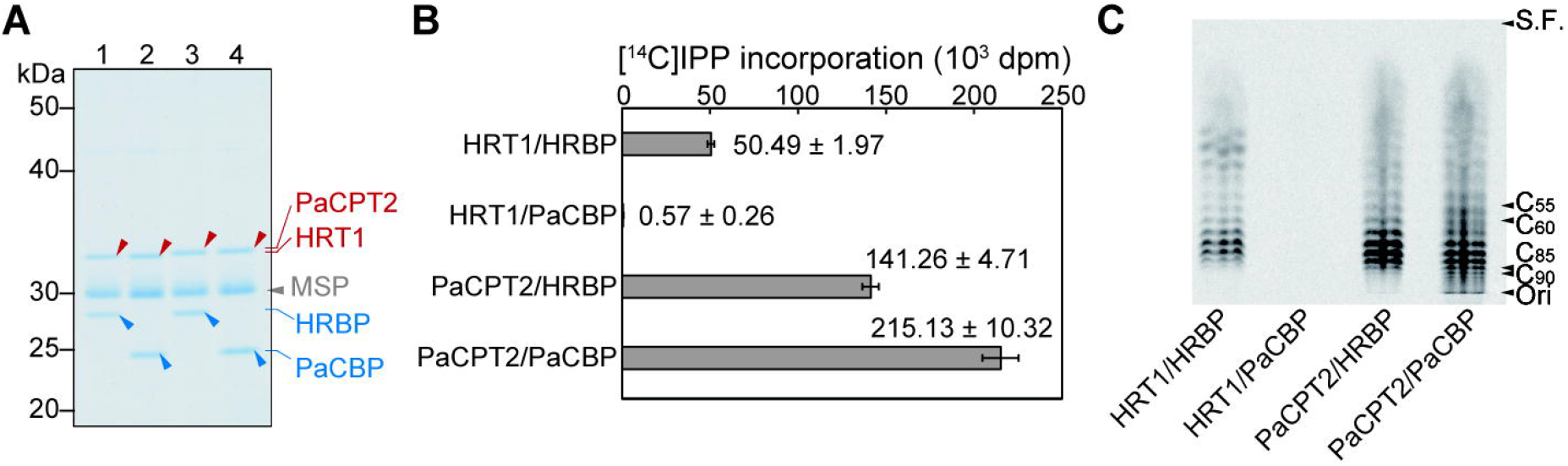
Prenyltransferase subunit compatibility between *H. brasiliensis* and *P. argentatum*. (**A**) Protein-nanodisc complexes were prepared by co-translation of HRT1 and HRBP (lane 1), HRT1 and PaCBP (lane 2), PaCPT2 and HRBP (lane 3), or PaCPT2 and PaCBP (lane 4). The purified complexes were analyzed by SDS-PAGE and CBB staining. Arrowheads indicate HRT1 or PaCPT2 (red), HRBP or PaCBP (blue), or MSP (gray). (**B**) Polyisoprene synthesis activity assay for the purified complexes in (**A**). The incorporation of [^14^C]IPP into macromolecules extracted with 1-butanol was measured. Data are means ± SD from three independent experiments. (**C**) Analysis of chain length for the ^14^C-labeled polyisoprenes extracted by 1-butanol was performed by TLC and autoradiography.

## 4. Discussion

We here demonstrated that co-expression of HRT1 and HRBP of *H. brasiliensis* in the presence of nanodiscs reconstitutes prenyltransferase activity in vitro. As far as we are aware, this is the first demonstration that the combination of these proteins gives rise to such activity on a lipid-bilayer membrane. The order of sequential expression of HRT1 and HRBP did not affect enzyme activity, suggesting that each single subunit associated with nanodiscs is stable during purification and readily accepts the other subunit to form a functional prenyltransferase. We also found that *P. argentatum* homologs of these *H. brasiliensis* proteins also reconstituted prenyltransferase activity on nanodiscs. The prenyltransferase subunits of the two species were only partially compatible, with full activity being achieved only with the subunit pairs from the same species. These results suggest that the distantly related plants *H. brasiliensis* and *P. argentatum* deploy similar heterosubunit enzymes for the biosynthesis of polyisoprenes.

On the other hand, the reconstituted prenyltransferases in the present study did not generate rubber-size polyisoprenes, possibly reflecting the technical difficulty of preparing a monolayer membrane that mimics rubber particles. Other proteins, such as REF and SRPP, together with a more suitable membrane system, may thus be required to support the activity of the prenyltransferase core enzyme in rubber-producing plants [6,20,21].

## Abbreviations

BSA: bovine serum albumin
CBB: Coomassie brilliant blue
CBP: CPT binding protein
CPT: *cis*-prenyltransferase
HRBP: HRT1-REF bridging protein
HRT: *Hevea brasiliensis* rubber transferase
IMAC: immobilized metal affinity chromatography
IPP: isopentenyl pyrophosphate
MSP: membrane scaffold protein
NR: natural rubber
PAGE: polyacrylamide gel electrophoresis
REF: rubber elongation factor
SRPP: small rubber particle protein
TLC: thin-layer chromatography.

## Declaration of competing interest

We declare no conflict of interest related to this study.

## Acknowledgments

We thank Frank Bernhardt (Goethe University) for suggestions regarding preparation of nanodiscs. This work was supported in part by a Grant-in-Aid from the Japan Society for the Promotion of Science (21H02115 to S.Y. and Y. Tozawa)

## Appendix A. Supplementary data

Supplementary data related to this article can be found online.

## Notes

### Competing Interest Statement

The authors have declared no competing interest.

