## Supplementary materials for "Reconstitution of prenyltransferase activity on nanodiscs by components of the rubber synthesis machinery of the Para rubber tree and guayule"

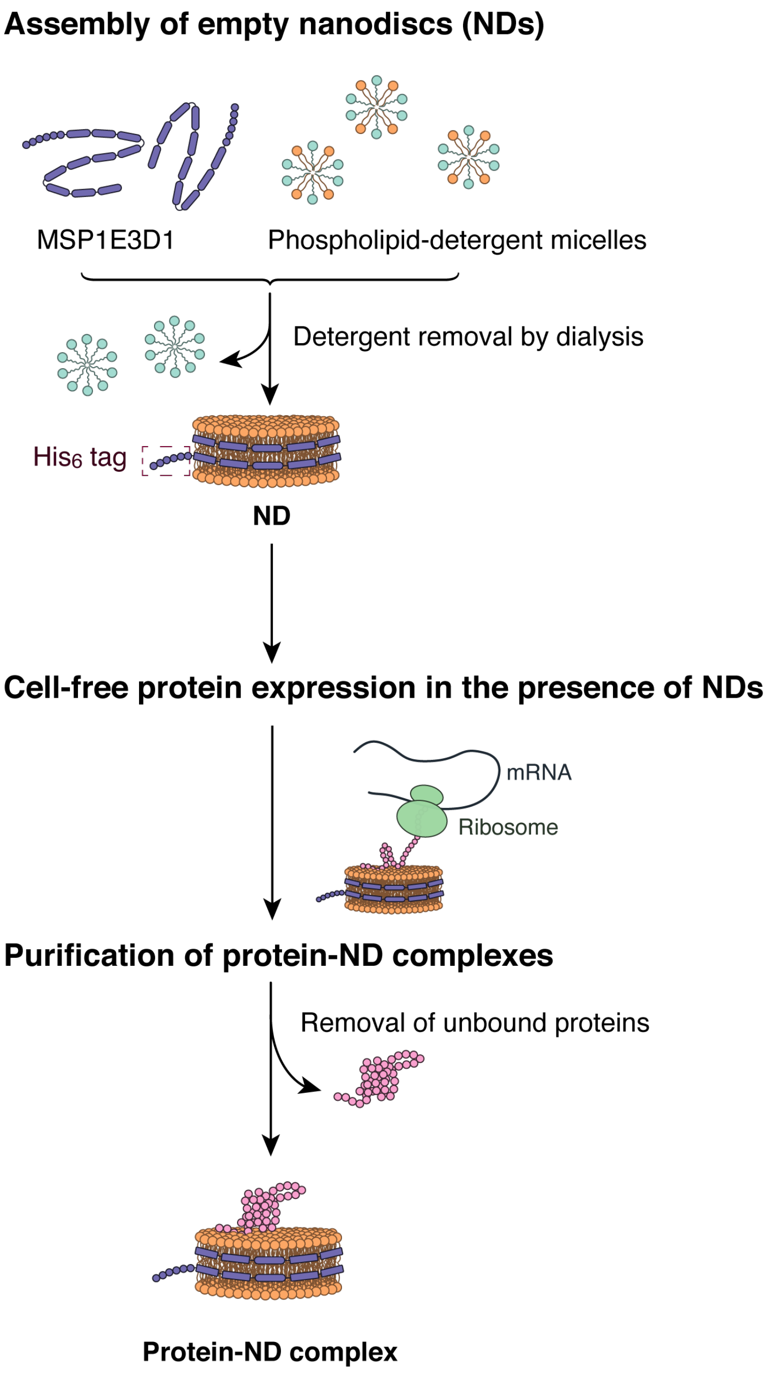


**Supplementary Fig. S1.** Procedure for preparation of protein-nanodisc complexes. Empty nanodiscs (NDs) were prepared from purified membrane scaffold protein (MSP1E3D1) and asolectin, and were then purified by IMAC in a manner dependent on the $NH_2$-terminal $His_6$ tag of MSP. The purified nanodiscs were then added to a cell-free translation system, and co-translationally assembled (or bound) proteins were co-purified with the nanodiscs by IMAC.

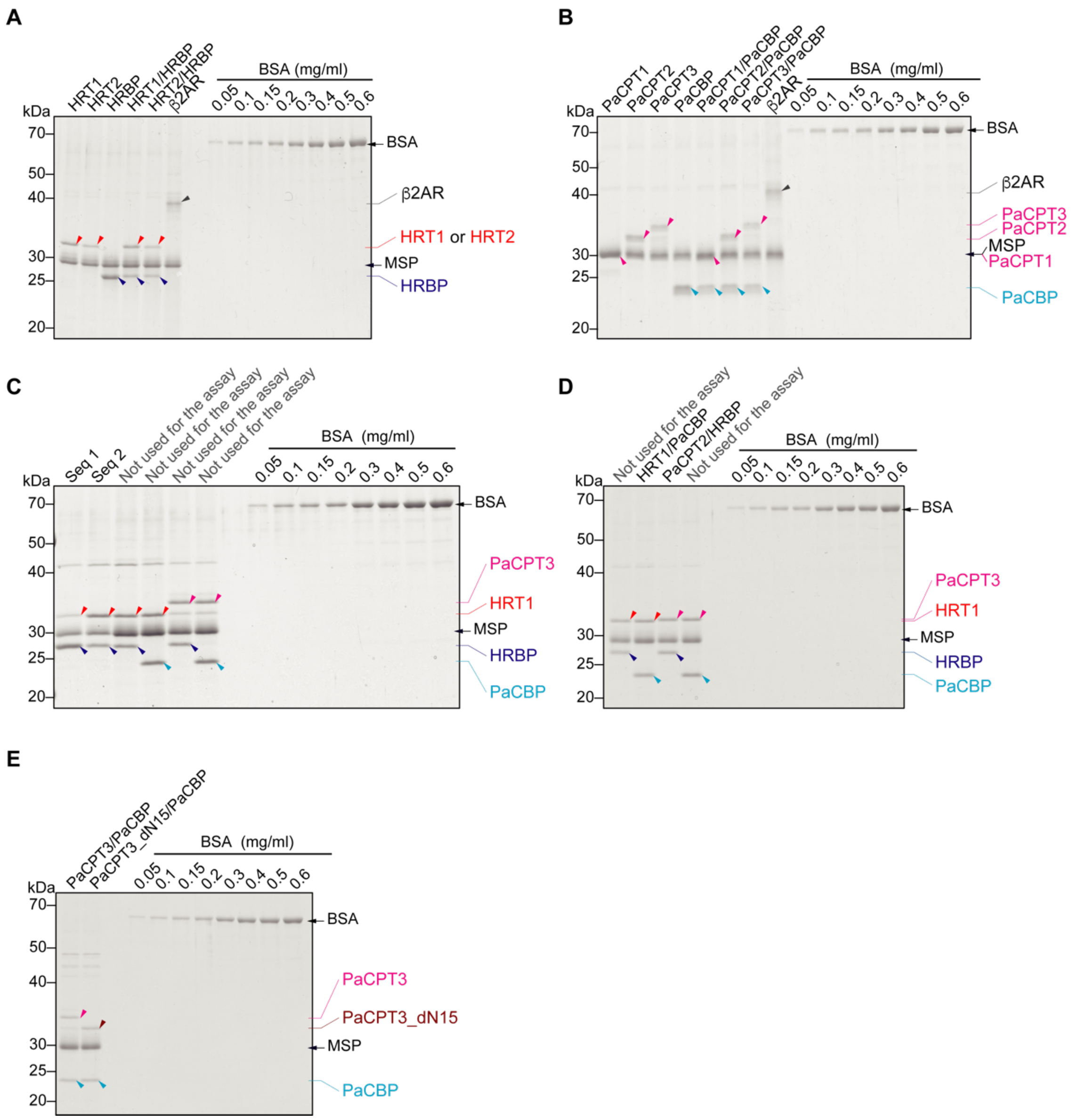


**Supplementary Fig. S2.** Estimation of protein concentration for protein-nanodisc complexes by SDS-PAGE and CBB staining and comparison with a BSA standard. *Hevea brasiliensis* (HRT1, HRT2, and HRBP) and *P. argentatum* (PaCPT1, PaCPT2, PaCPT3, PaCPT3_dN15, and PaCBP) proteins as well as human β2AR were assembled with asolectin nanodiscs by cell-free protein synthesis with a wheat-germ translation system. The resulting protein-nanodisc complexes were purified by Ni-NTA column chromatography and then subjected to SDS-PAGE and CBB staining together with BSA standard solutions. The protein concentration of each sample was estimated from band intensity with ImageJ software. The samples shown here include those used for prenyltransferase activity assays in Figure 1 (**A**), Figure 3 (**B**), Figure 2 (**C**), Figure 4 (**D**), and Figure S4 (**E**). Some of the samples were not used for the enzyme assay.

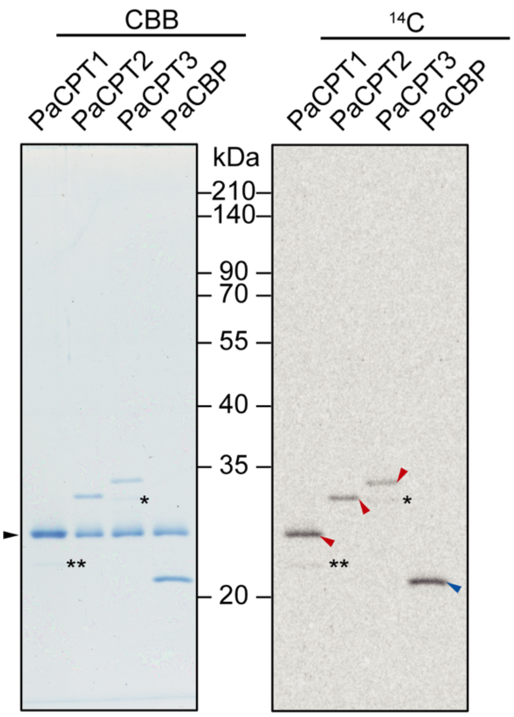


**Supplementary Fig. S3.** Analysis of additional products in cell-free translation reactions. PaCPT1, PaCPT2, PaCPT3, and PaCBP were synthesized in a cell-free system in the presence of nanodiscs and [$^{14}$C]leucine. The purified protein-nanodisc complexes were then subjected to SDS-PAGE, and the gel was stained with CBB (left panel) and subjected to autoradiography (right panel). Single and double asterisks indicate minor products in the PaCPT3 and PaCPT1 reaction mixtures, respectively. The black arrowhead indicates the band for MSP of the nanodiscs, which overlaps with that for PaCPT1. Red and blue arrowheads indicate the major $^{14}$C-labeled proteins, whose apparent molecular sizes reflect their predicted molecular masses of 31.3 kDa for PaCPT1, 33.2 kDa for PaCPT2, and 34.8 kDa for PaCPT3 and of 28.7 kDa for PaCBP, respectively.

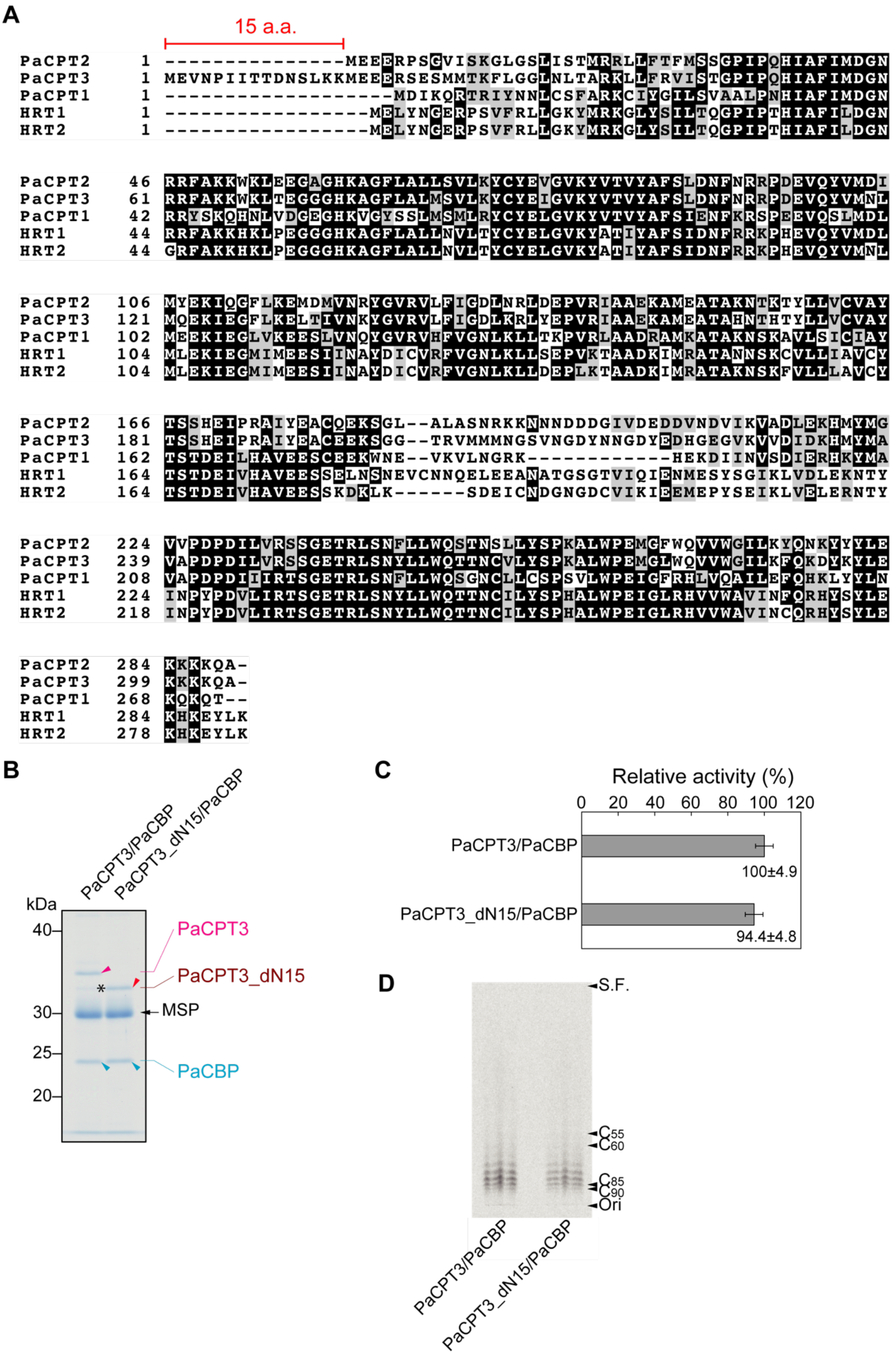


**Supplementary Fig. S4.** Analysis of a truncated form of PaCPT3. (**A**) Alignment of the predicted amino acid sequences of HRT1, HRT2, PaCPT1, PaCPT2, and PaCPT3. Hyphens indicate gaps introduced to optimize alignment. Identical or similar residues among the five proteins are indicated by dark and light shading, respectively, and residue numbers are shown on the left of the sequences. PaCPT2 and PaCPT3 are closely related, sharing the highest sequence identity (89%) among the five sequences. PaCPT3 possesses a methionine at position 16 that corresponds to the first methionine of PaCPT2. (**B**) PaCBP as well as either a truncated form of PaCPT3 lacking the $NH_2$-terminal 15 amino acids (PaCPT3_dN15) or the wild-type protein were synthesized in a cell-free system in the presence of nanodiscs, and the purified protein-nanodisc complexes were characterized by SDS-PAGE and CBB staining. The asterisk indicates a protein in the PaCPT3 reaction mixture with a mobility similar to that of PaCPT3_dN15. (**C**) Relative prenyltransferase activity of PaCPT3/PaCBP-nanodisc and PaCPT3_dN15/PaCBP-nanodisc complexes. The raw data used to calculate relative activity are 22,683 ± 1105 dpm for PaCPT3/PaCBP and 21,406 ± 1096 dpm for PaCPT3_dN15/PaCBP (means ± SD from three independent experiments). (**D**) TLC analysis of extracts prepared from enzyme assay mixtures as in (**C**) with 1-butanol. Both of the PaCPT3/PaCBP and PaCPT3_dN15/PaCBP complexes showed similar patterns of polyisoprene chain length.
